# Accessory genomic epidemiology of co-circulating *Acinetobacter baumannii* clones

**DOI:** 10.1101/2021.03.26.436874

**Authors:** Valeria Mateo-Estrada, José Luis Fernández-Vázquez, Julia Moreno-Manjón, Ismael L. Hernández-González, Eduardo Rodríguez-Noriega, Rayo Morfín-Otero, María Dolores Alcántar-Curiel, Santiago Castillo-Ramírez

**Affiliations:** Programa de Genómica Evolutiva, Centro de Ciencias Genómicas, Universidad Nacional Autónoma de México, Cuernavaca, México; Laboratorio de Infectología, Microbiología e Inmunología Clínicas, Unidad de Investigación en Medicina Experimental, Facultad de Medicina, Universidad Nacional Autónoma de México, Ciudad de México, México; Hospital Civil de Guadalajara Fray Antonio Alcalde e Instituto de Patología Infecciosa y Experimental, Centro Universitario de Ciencias de la Salud, Universidad de Guadalajara, Guadalajara, Jalisco, México; Laboratorio de Bacteriología Médica, Posgrado en Ciencias Quimicobiológicas, Escuela Nacional de Ciencias Biológicas, Instituto Politécnico Nacional, Carpio y Plan de Ayala SN, Ciudad de México, México

**Keywords:** Genomic epidemiology, bacterial clones, transmission dynamics, accessory genome, *Acinetobacter baumannii*, molecular epidemiology

## Abstract

*A. baumannii* has become one of the most important multidrug resistant nosocomial pathogens all over the world. Nonetheless, very little is known about the diversity of *A. baumannii* lineages co-existing in hospital settings. Here, using whole-genome sequencing, epidemiological data and antimicrobial susceptibility tests, we uncover the transmission dynamics of extensive and multidrug resistant *A. baumannii* in a tertiary hospital for a decade. Our core genome phylogeny of almost 300 genomes suggests that there were several introductions of lineages from international clone 2 into the hospital. The molecular dating analysis shows that these introductions happened between 2004 and 2015. Furthermore, using the accessory genome, we show that these lineages were extensively disseminated across many wards in the hospital. Our results demonstrate that accessory genome variation can be a very powerful tool for conducting genomic epidemiology. We anticipate future studies employing the accessory genome as a phylogenomic marker over very short microevolutionary scales.

## INTRODUCTION

Antimicrobial drug resistance is one of the most important health issues worldwide. The ESKAPE (*Enterococcus faecium*, *Staphylococcus aureus*, *Klebsiella pneumoniae*, *Acinetobacter baumannii*, *Pseudomonas aeruginosa*, and *Enterobacter* species) group have many acquired antimicrobial resistance genes (ARGs) and are a major source of deadly infections all over the world ^1^. The World Health Organization list of bacterial pathogens urgently requiring novel antibiotics ranked *A. baumannii* at the highest priority status (priority 1: critical) ^2^ Importantly, many *A. baumannii* nosocomial infections are due to isolates resistant to at least one agent in three or more antimicrobial agents, namely multidrug resistant (MDR) phenotypes, which make them very difficult to treat. Clearly, *A. baumannii* is a global public health issue that needs to be seriously considered.

Over the last decades whole-genome sequencing (WGS) has become the ultimate approach to study the dissemination of many bacterial pathogens ^3–7^ WGS has been extremely useful in the case of *A. baumannii*, where the very dynamic nature of its genome renders multilocus sequence typing (MLST) faulty ^8^ WGS has been used to study the spread of clones of *A. baumannii* at national and even at continental levels ^9,10^. However, unlike other important bacterial pathogens, WGS has hardly been used to analyse clone diversity within hospital settings. Actually, very little is known about whether one or several different lineages are the cause for the MDR infections found in hospital settings, not least in Latin America. Nonetheless, a recent study has suggested that different lineages could be co-circulating in the National Institute of Oncology in Mexico City ^9^.

When using WGS for epidemiological investigations, the most common approach is to use core Single Nucleotide Polymorphisms (SNPs) phylogenies or SNPs differences between isolates. This approach has resulted effective in most cases. However, considering very short periods of time, there are instances where hardly any SNPs have had accrued between strains, not least in the core genome. For instance, in a recent study ^11^ that analysed 24 *A. baumannii* strains collected from the blood of a single patient (over a period of 6 months) some strains (sampled during the same month) were identical *as per* core SNPs. However, a few years ago, in a population genomics study we showed that acquisition and depletion of genes occurs way faster than the accumulation of mutations in *A. baumannii* ^12^. Therefore, this gene content variability can be very valuable for studying the evolutionary relationships of isolates at very short periods of time.

The goal of our study was to used WGS, along with epidemiological data and antimicrobial susceptibility testing, to analyse the diversity of clones (and their antimicrobial resistance patterns) in a hospital sampled for a decade. To that end, we carried out a genomic epidemiology study to characterize the lineages of *A. baumannii* in a tertiary, teaching hospital in Guadalajara (Hospital Civil de Guadalajara [HCG]), Mexico. Notably, besides the core genome, we also used the accessory genome to establish the dissemination of lineages across different wards.

## RESULTS

### Extensive and multidrug drug resistant lineages from International Clone 2

We sequenced 73 isolates of *A. baumannii* sampled between 2007 and 2017 from the HCG (see methods and Supplementary Table 1). The HCG is a tertiary, teaching hospital in Guadalajara, Jalisco (Western Mexico), and it has 899 beds. All these isolates belonged to the pulse-type 22, which in previous studies ^13,14^ was the most prevalent clone in this hospital. We used the genome sequences to conduct Sequence Type (ST) assignation, considering the Oxford MLST scheme ^15^. These isolates belonged to just four STs (see Figure 1). ST417 was the most frequent with 38 isolates, followed by ST208 having 23 isolates; whereas ST136 and ST369 had 7 and 5 isolates, respectively. Of note, all these STs are part of the international clone 2 (IC2). We conducted antimicrobial susceptibility tests on the isolates (see methods).

**Figure 1.**
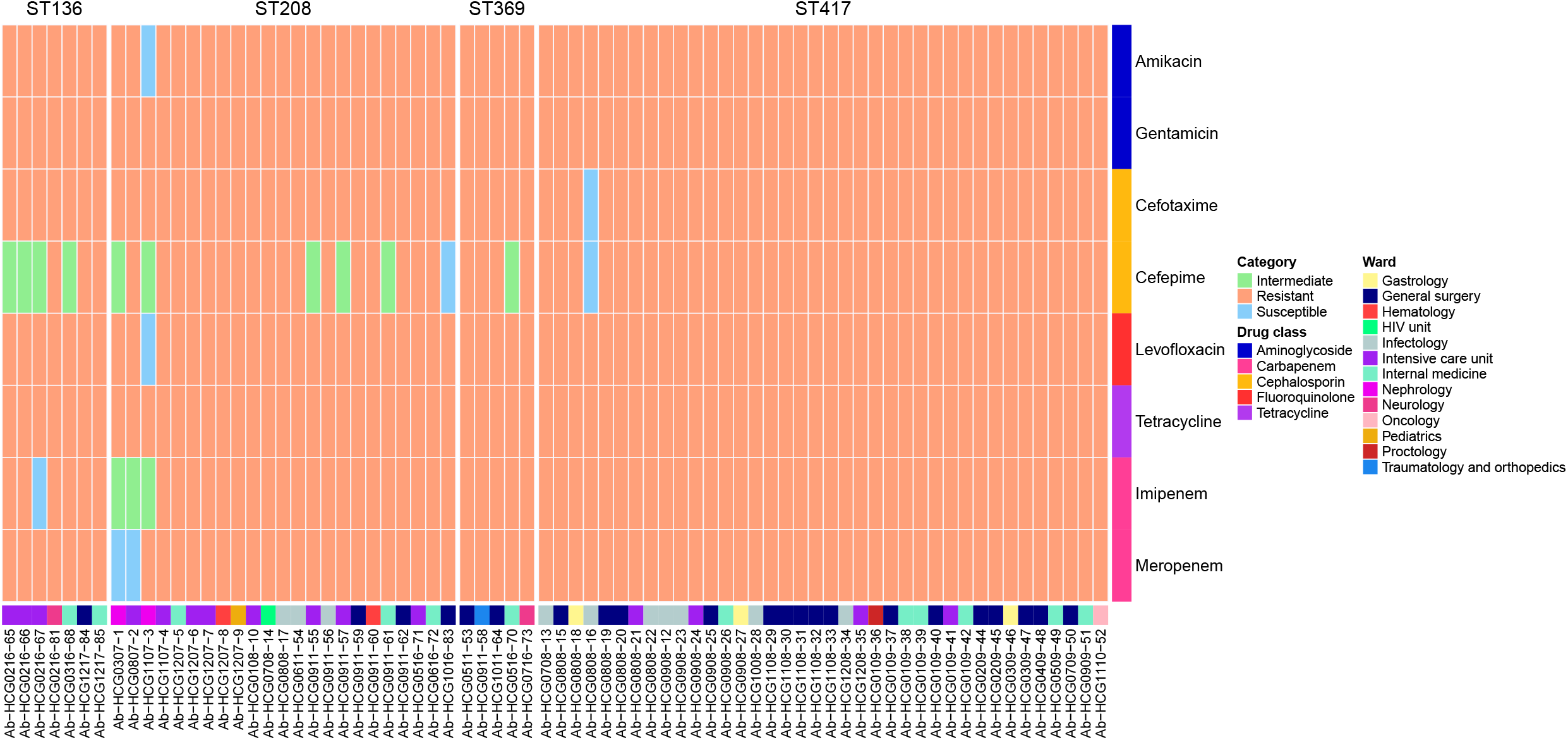
Antibiotic resistance patterns of the 73 *A. baumannii* isolates from the HCG. Different drug classes were tested and the actual MICs for each isolate, and each drug, are provided in supplementary table 2. For each isolate we also provide the ward from which it was sampled. The ST assignation for the isolates is at top. Re the strain names, the numbers after the “HCG” initials give the month (first two digits) and the year (third and fourth digits) of sampling. For instance, Ab-HCG**0216**-65 was sampled in February 2016.

Supplementary Table 2 gives the minimum inhibitory concentration (MIC) values for each isolate and each drug tested. Remarkably, all the isolates were resistant to many antibiotics (see Figure 1). We found some variation across the different STs. While ST136 and ST208 had less resistance, ST369 and ST417 had more resistance. All but one of the ST369 isolates were resistant to all the antibiotics tested; whereas for the ST417 isolates also all but one were resistant to all the antibiotics tested. Most of the isolates, irrespective of their ST assignation, were extensively drug-resistant (XDR) (resistant to at least one agent in all but two or fewer antimicrobial categories). There were a few isolates that were MDR isolates within ST208. These results suggest that these isolates, regardless of which ST they belong to, are either MDR or XDR. We cannot say that there are pan-drug resistant (PDR) isolates (resistant to all agents in all antimicrobial categories) since we did not test all the antibiotics. However, we are safe to say that the vast majority of the isolates are extensive drug-resistant and very likely some of these may be PDR. Taken together, these results show that different MDR and XDR lineages (STs), all assigned to IC2, were found in the hospital.

### Several introductions into the hospital between 2004 and 2015

To have a better idea of the introduction of these STs into the hospital, we downloaded publicly available genomes showing the same STs as the HCG isolates. Then, considering the HCG isolates and the publicly available genomes, we constructed a Maximum Likelihood core genome phylogeny (see methods). This phylogeny had 299 genomes and is shown in Figure 2. The phylogeny clearly shows that the STs were introduced in independent events into the hospital (see stars in Figure 2). The HCG isolates of the most frequent ST, namely ST417, all clustered together. Given the monophyletic nature of this ST in the hospital, one single introduction of this STs into the hospital is the most plausible explanation. In the case of HCG ST208 isolates, there were two introduction events. We noted that all the isolates but isolate Ab-HCG1107-3 cluster together, implying one single introduction event. Whereas Ab-HCG1107-3, located on a distant branch, was introduced independently. Considering the HCG ST136 isolates, they form a paraphyletic group as two isolates (Ab-HCG0516-70 and Ab-HCG0716-73) assigned to the ST369 clustered with them. However, all the HCG ST136 isolates plus the two ST369 isolates mention above were introduced in one single even into the hospital. Finally, another three ST369 isolates (Ab-HCG0511-53, Ab-HCG0911-58, Ab-HCG1011-64) were introduced in another event. Then, we wanted to know the time of introduction of these STs into the hospital. Thus, we conducted a molecular dating analysis (see methods). We only estimated the time to the Most Recent Common Ancestor (tMRCA) for ST417, ST208 (not including the isolate introduced in the independent event), and ST136. Figure 3 shows the marginal posterior densities of the tMRCAs for the three STs considered. From this figure is clear that ST208 and ST417 were introduced very closed to one another; whereas ST136 was introduced a few years later. ST208 was the first lineage to get to the hospital and it was introduced in early 2006, 95 % High Posterior Density interval (HPD), mid 2004 to late 2006. ST417 was introduced about one year later, in early 2007 (95 % HPD, late 2005 to late 2007). Some five years later the ST136 was introduced (tMRCA 2013; 95 % HPD, early 2011 to mid 2015). Taken into account the 95 % HPDs for the three STs, these lineages were introduced into the hospital between 2004 and 2015. Collectively, these data suggest that five events account for the introduction of these isolates into the HCG and these introductions occurred more or less over a decade.

**Figure 2.**
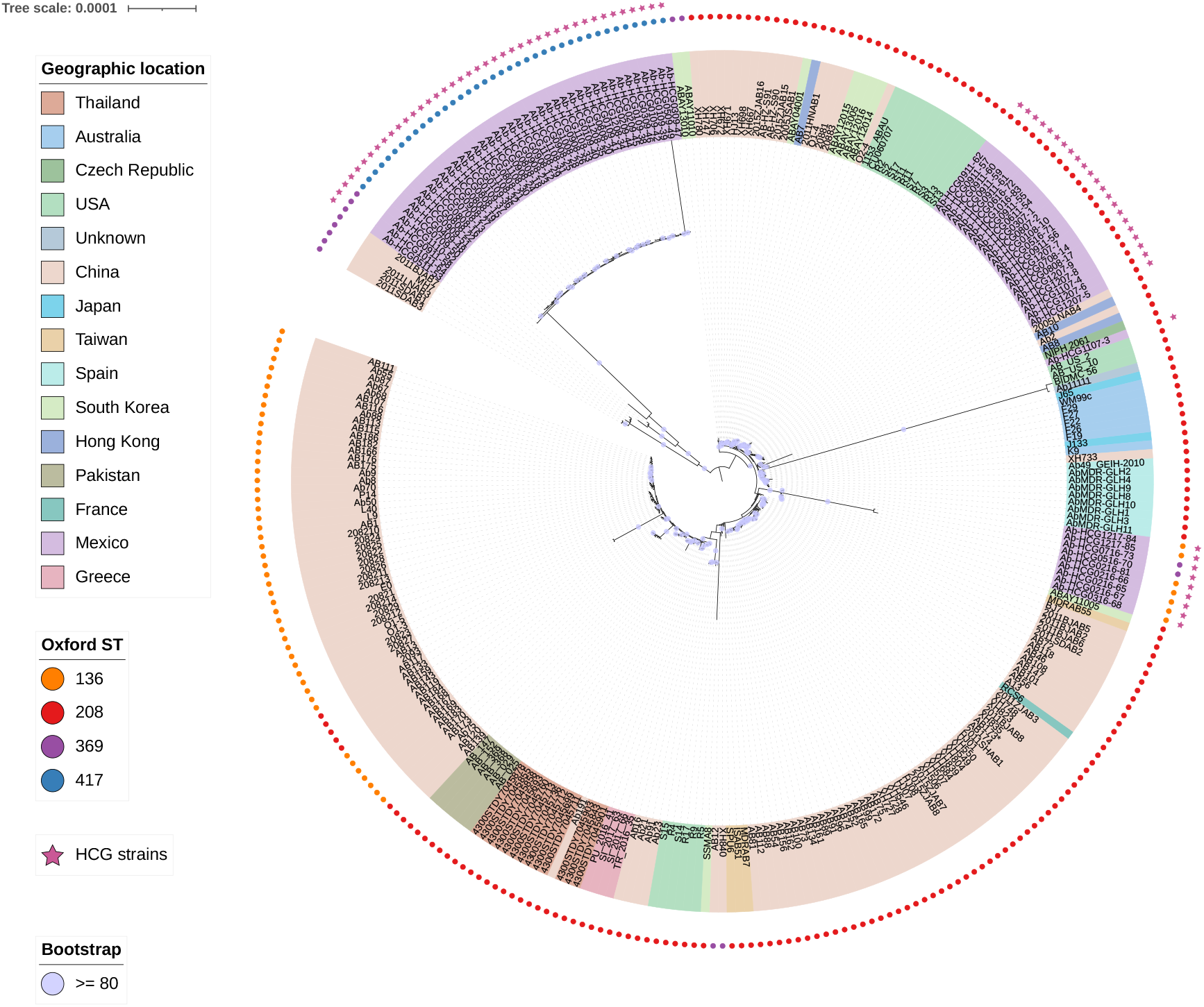
Phylogeny of the 73 *A. baumannii* isolates from the HCG plus publicly available genomes. The phylogeny is annotated with the geographic region and the ST. The stars mark the isolates from the HCG. Bootstrap values equal to or higher than 80 are shown with violet dots at the internal nodes of the tree. The scale bar gives the number of substitutions per site.

**Figure 3.**
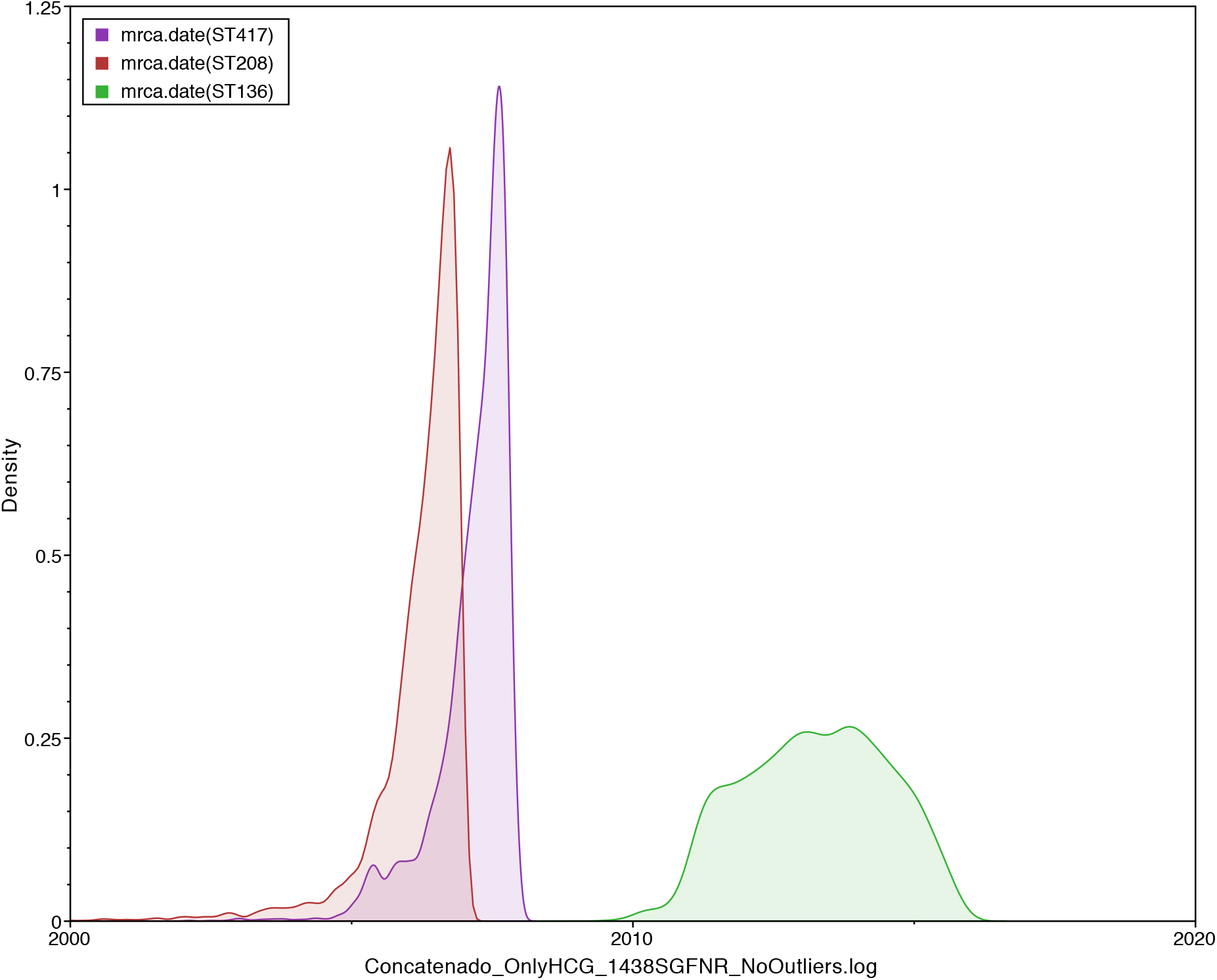
Molecular dating analysis of the main ST. Marginal posterior density for the tMRCA for ST136 (green), ST208 (red), and ST417 (purple). The x-axis shows the time between 2000 and 2020.

### High transmission across wards

The isolates sequenced were collected from 13 different wards in the hospital. Figure 1 shows the distribution of isolates per ward and year. Intensive Care Unit (ICU) and General Surgery were the most represented wards among the isolates sequenced. The distribution of isolates per ward suggests that the different STs have spread to many wards in the hospital. To further analyze this, we conducted two analyses just comparing isolates from the same ST. In each analysis, we compared isolates between and within wards and counted the number of differences. In the first analysis, we counted the number of core SNPs telling apart isolates, whereas in the second analysis we counted the number of genes differing between isolates (see Figure 4). These analyses were only conducted for ST417 and ST208, the 2 most frequent STs. If isolates were mainly disseminating in the same ward, one would expect considerably less differences (either core SNPs or number of genes) when comparing isolates from within the same ward than when comparing isolates from different wards. On the contrary, if there were high transmission across wards, the number of differences when comparing intra versus inter ward isolates would not be that different. Our analyses supported the latter scenario, whether we considered core SNPs (upper panels) or the number of genes differing between isolates (lower panels). Considering both measures, the distribution of intra ward comparisons overlaps with the distribution of inter ward distribution (see Figure 4). We did not find significant differences comparing the distributions of core SNPs (Wilcoxon–Mann–Whitney test, ST417 p-value = 0.6872 and ST208 p-value = 0.7834) nor comparing the distribution of different genes (Wilcoxon-Mann-Whitney test, ST417 p-value = 0.6568 and ST208 p-value = 0.6478).

**Figure 4.**
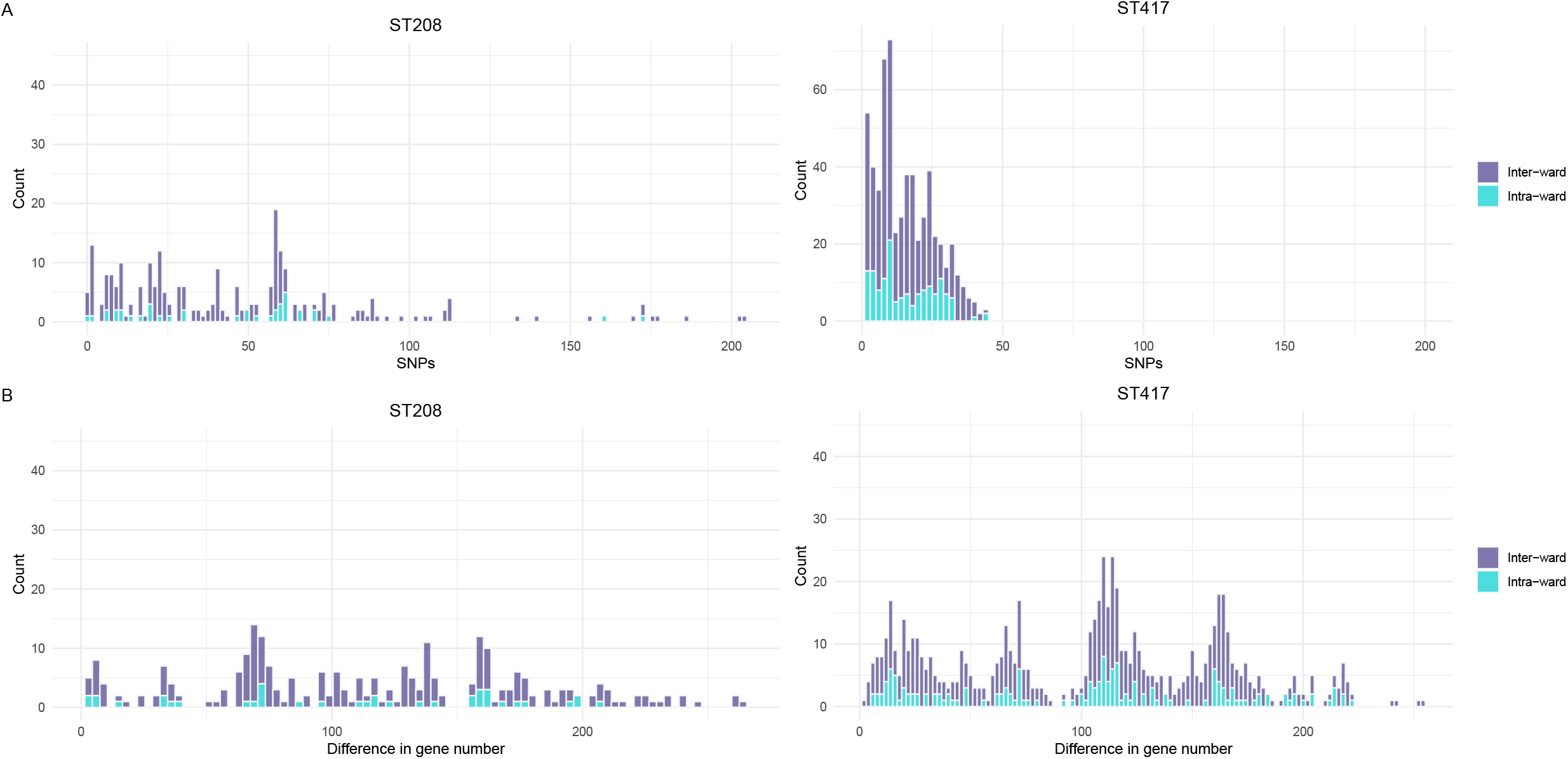
Inter and intra ward transmission analysis. We conducted pairwise comparisons for the isolates of ST208 (left-hand side) and ST417 (right-hand side), which were the most abundant STs. Comparisons considering isolates from the same ward are shown in cyan (light blue) bars, whereas comparison of isolates from different wards are the purple bars. The top panels give the differences in terms of core SNPs, whereas the bottom panels show the differences in gene content.

To further explore the high transmission of isolates across wards, we also analyzed the gene content variation among the isolates, which according to our previous data accrues faster than *de novo* mutations in *A. baumannii*. To this end, we constructed a gene correlation matrix for the 73 isolates (see Figure 5). In this figure one can see regions (squares) of dark blue, which denote isolates with very similar gene content and, therefore, very closely related to one another. We noted that these regions are composed of isolates from different wards (see vector “Wards” at the bottom of the figure). This pattern applies to all the four STs. For instance, the isolates Ab-HCG0908-26, Ab-HCG0908-21, Ab-HCG0908-24 and Ab-HCG0908-33 (all from ST417) are very similar in terms of their gene content yet they were isolated from three different wards (ICU, internal medicine, and general surgery). Considering ST208, isolates HCG1207-8, HCG1107-4, HCG1207-7, HCG1207-5, HCG1207-6, were isolated from ICU, internal medicine, and hematology, and again have a pretty similar gene content. Then, to compare the discriminative typing power of the core genome and the accessory genome, we constructed another two trees just including the 73 HCG isolates. One tree was constructed using the core genome and the second tree was based on a gene content distance matrix (see methods). These two trees are presented in Figure 6. Both trees reinforce the idea of high transmission across wards, as isolates for any given ward very often had their closest related isolate coming from a different ward. Of note, the core genome tree (right-hand panel Figure 6) had several polytomies and these affect the different STs, ST417 and ST208 being the most outstanding cases. Thus, some isolates were identical to other isolates under the core genome tree. On the contrary, the phylogenetic tree based on gene content (left-hand panel Figure 6) showed far greater resolution. In this tree all the isolates could be differentiated from one another. Therefore, gene content variation (accessory genome) had more discriminative genotyping resolution than the core genome. All in all, these analyses demonstrate that there is frequent transmission of isolates across many different wards in the hospital. Furthermore, the analyses of the accessory genome were of paramount importance to describe the transmission dynamics of these isolates.

**Figure 5.**
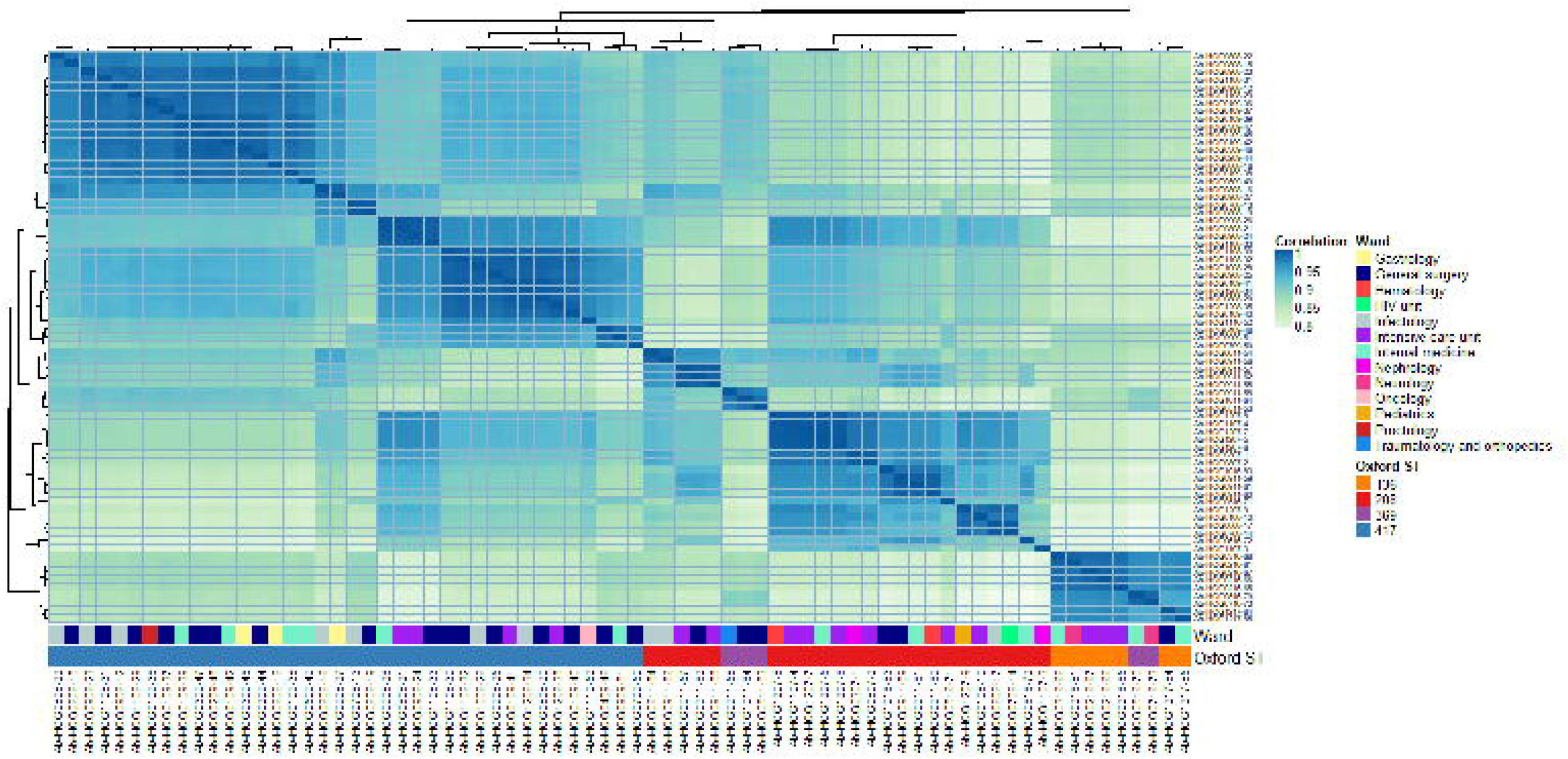
Gene content variation among the isolates from the HCG. Heat map visualising the gene content correlation matrix. The heat map is annotated with the ward and the ST for each isolate. The top and side dendrograms show the clustering by similarity in gene content. The darker the blue, the more similar the isolates are in gene content.

**Figure 6.**
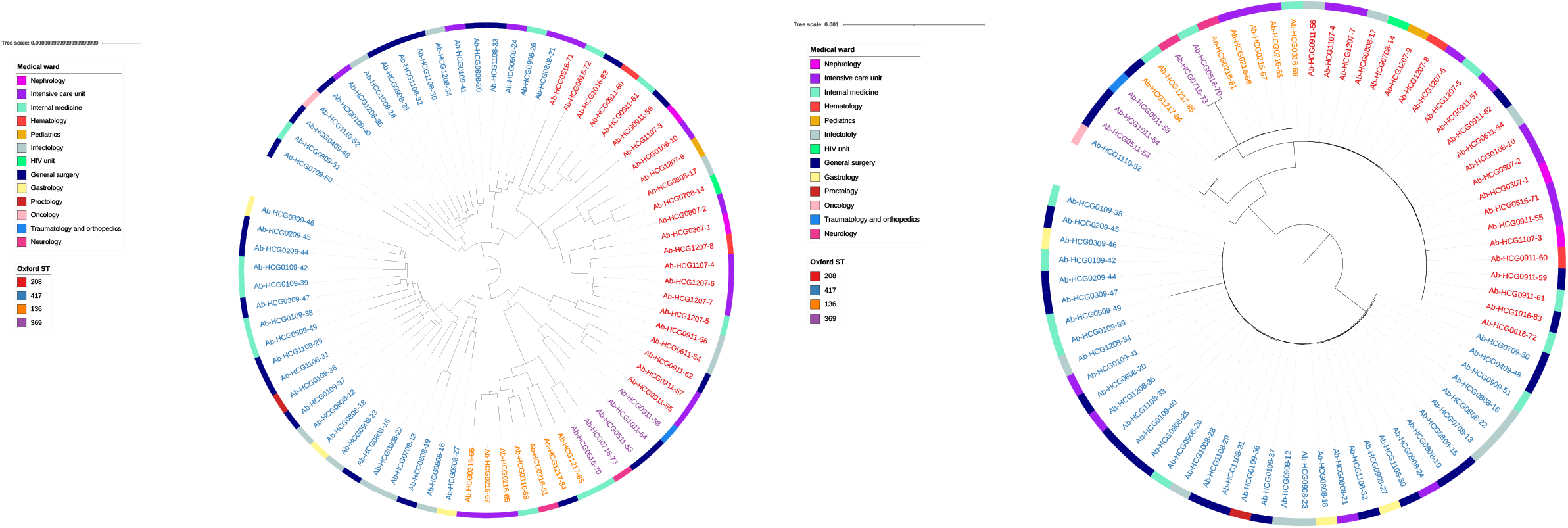
Accessory and core genome trees for the HGC isolates. Both trees were constructed only considering the newly sequenced 73 *A. baumannii* isolates from the HCG. Left-hand side panel, Neighbor-joining tree based on a gene content distance matrix. Right-hand side panel, ML tree based on the alignment used for the ML phylogeny in Figure 2, the alignment was edited to only contain the 73 isolates from the HCG. The isolates are color-coded *as per* ST, whereas the external ring provides the wards for the isolates.

## DISCUSSION

Over the last decade WGS has been successfully used to determine transmission dynamics at very different levels; from local outbreaks to global spread of pathogens. However, WGS has been hardly used to study different lineages of *A. baumannii* in a single setting. In this study, we used both core and accessory genome to get an uber-resolution of the transmission dynamics of co-circulating lineages in a tertiary hospital over a decade. We found 4 lineages (ST136, ST208, ST369, and ST417) and all were from the IC2. This clone has been described in many parts of the world. However, this clone has not been described as one of the most relevant clones in Mexico in recent genomic studies ^9,12^. Nonetheless, IC2 has been described in other countries in Latin America ^16^ or the United States ^17^, which shares an extensive border with Mexico. Our phylogenetic analysis based on the core genome indicates that these STs were introduced in different events into the HCG. These introduction events occurred in 2006 and 2007, and in 2013. Clinically speaking, it is relevant that the isolates from these STs were either MDR or even XDR. Also, most of them were resistant to carbapenems. Contrary to the idea of just having one predominant clone at any time, we found different STs in the period of time this study encompasses. Hence, it seems that different lineages can be co-existing within the same hospital setting.

To properly analyze ward dissemination, we used both core and accessory genome. Although the accessory genome analysis had way more resolution, both analyses clearly showed that the STs have been extensively disseminated in many wards. Thus, there is no population structuring according to wards. From a practical point of view, these results suggest that infection control measures were not able to contain the dissemination of these STs in the hospital. In accordance with our previous work ^8^, we noted that several STs (ST208, ST369 and ST136) have problems (*i.e*., they do not form monophyletic groups) properly genotyping the isolates assigned to them. This highlights the need for the use of more powerful genotyping approaches for *Acinetobacter baumannii*. Clearly, if possible, WGS should be the method of choice. Over the last decade most bacterial genomic epidemiology studies have used core SNPs to reconstruct the transmission dynamics of pathogens. This is on account of the fact that *de novo* mutations accrue on the same time scale (ideally, faster) than the transmission events inferred. However, for some bacterial species, *A. baumannii* being the case in point, this might not happen at very short time scales. In our study, to overcome this, we used gene content variation, which occurs faster than mutations ^12^, to have a higher genotyping resolution at very short time scales. Remarkably, our results show that this type of alternative genomic variation can be used to track the transmission events at very short periods of time. Ideally, one can use both core and accessory genome to conduct genomic epidemiology at an unprecedented level; going beyond what the core genome variation can resolve by itself. We think the use of the accessory genome (as an extra phylogenomic marker) can be particularly useful for epidemiological investigations of outbreaks.

In conclusion, our study shows that both the accessory and the core genome can be very powerful tools to understand the transmission dynamics of bacterial pathogens. We anticipate our study to be a reference point for more elaborate analyses using the accessory genome as a very useful phylogenomic marker over very short microevolutionary scales.

## MATERIAL AND METHODS

### DNA extraction, genome sequencing and antibiotic susceptibility testing

The initial collection had 134 isolates of *A. baumannii*, which were identified in previous studies ^13,14^, and these isolates covered a decade, from 2007 to 2017. Several pulse types were identified among these 134 isolates but pulse-type 22 was the most frequent by far. We sequenced 76 isolates (all pulse-type 22) trying to represent as much as possible the proportion of isolates per year in the initial collection. Three of them had to be excluded due to quality issues; our final data set has 73 genomes. Supplementary Table 1 provides the details (date, ST, ward, *etc*.) of these genomes. Genomic DNA was extracted using the QIAamp DNA Mini Kit (Qiagen, Hilden, Germany) according to the manufacturer’s instructions from a single colony grown in LB medium grown overnight at 37°C ^18^. The genome sequencing of the isolates was carried out at the Instituto Nacional de Medicina Genómica (https://www.inmegen.gob.mx/) in Mexico City. A 2 × 250-bp configuration, employing an Illumina Miseq platform, was employed. We trimmed the first and last five bases in each read and the poor-quality bases with Trim Galore (https://github.com/FelixKrueger/TrimGalore). We assembled the genomes with SPAdes ^19^ (see Supplementary Table 1 for assembly statistics) and then we annotated and checked the contamination and completeness of each genome as in ^20^. We used the PubMLST database ^21^ to assign Sequence Types, under the Oxford MLST scheme, to the newly sequenced isolates. We also downloaded over 200 genomes from the NCBI that had the same STs as those assigned to the newly sequenced isolates (see Supplementary Table 1). The MIC was performed for the 73 isolates sequenced using serial dilution on agar following the guidelines of the Clinical and Laboratory Standards Institute (CLSI) ^22^. The antimicrobial agents tested against the 73 isolates include amikacin (AMK), gentamicin (GEN), cefotaxime (CTX), cefepime (FEP), levofloxacin (LVX), tetracycline (TET), imipenem (IPM) and meropenem (MEM). The MIC was determined as the lowest concentration of antibiotics in which the *A. baumannii* growth was inhibited. Supplementary Table 2 gives the MIC values for each isolate and each drug tested. The classification of each isolate as susceptible or resistant was established according to the CLSI breakpoints ^22^. *Escherichia coli* strain ATCC 25922 was employed for quality control tests.

### Phylogenies, recombination analysis and molecular dating

We constructed a Maximum Likelihood phylogeny on the concatenated alignment of single gene families without recombination, we proceeded as in a previous study ^9^. Single gene families were identified analysing the output of Roary ^23^ and we employed PhiPack ^24^ to identify recombination. We found 1654 singles gene families without recombination (SGFwR). The SGFwR were aligned and the phylogenetic tree was constructed using RAxML ^25^ as in ^20^. We annotated the phylogenetic tree with iTOL ^26^. We also ran another tree with this alignment but only considering the 73 isolates from the HCG. We implemented a molecular dating analysis using BEAST 2 ^27^. The analysis was run on the concatenated alignment of the SGFwR considering the 73 HCG isolates. We used TempEst ^28^ to check the temporal signal within the data and we excluded some of the isolates whose genetic divergence and sampling date were unusual. We found a strong temporal signal, the regression of dates of sampling against the root-to-tip distances was 0.9074. We used the TrN+G as the site model, for this was selected by jModelTest ^29^ as the best model under the Akaike information criterion. We used a log-normal relaxed molecular clock model, which was calibrated using the sampling dates of the isolates. The analysis was run for 900,000,000 generations and samples were taken every 900,000 generations. The analysis was run twice to make sure the results were consistent. In both analyses we ensured that the effective sampling size of the likelihood of the tree was well above 200.

### Comparisons intra versus inter ward, gene content correlation matrix, and gene content tree

To evaluate the dissemination across wards we employed two measures. One measure was the number of core SNPs between pairs of isolates. For this, we made use of all the SNPs found in the concatenated alignment of 2048 single gene families without recombination for only the 73 HCG isolates (we ran another pangenome analysis using Roary just with the HCG isolates). For this analysis, we discarded the isolates Ab-HCG0509-49 and Ab-HCG0611-54 which seem to be hypermutator strains. Whereas, the second measure counted the number of differing genes between pairs of isolates. For both measures we just compared the isolates from within the same ST and the pairwise comparisons were divided into intra and inter ward comparisons. We also constructed a gene content correlation matrix, as we have done previously in a couple of studies ^20^. From the pangenome analysis of the 73 isolates we produced a gene content matrix for all the sequenced HCG isolates. Then, using that matrix and applying the *cor()* function in R, a gene content correlation matrix was created. The matrix was visualized as heat map using the *ComplexHeatmap()* package ^30^ in R. To create a Neighbor-joining tree based on gene content, we created a gene content distance matrix (based on the gene content matrix) employing the *dist()* function in R and setting Euclidean distance. After that, the APE library ^31^ was used to construct a Neighbor-joining tree on the distance matrix.

## AUTHORS’ CONTRIBUTION

SCR conceived and supervised the study. VME conducted almost all the *in silico* analyses. SCR conducted the molecular dating analysis. MDAC supervised the DNA extraction and antibiotic susceptibility testing. JLFV carried out DNA extraction. JM conducted antibiotic susceptibility testing. MDAC, ERN, RMO provided the isolates and metadata. IGH downloaded the publicly available genomes and carried the ST assignation for them.

## ACKNOWLEDGEMENTS

We are grateful to Victor Manuel del Moral Chávez and Alfredo José Hernández Álvarez for their valuable job installing some of the bioinformatic programs used in this study. We also thank Joel Gómez Espíndola, Iván Uhthoff Aguilera, Maria Gabriela Guerrero Ruiz and Luis Fernando Lozano Aguirre Beltrán for technical support on several matters.

## SUPPLEMENTARY MATERIAL

**Supplementary Table 1**

List of the 299 *A. baumannii* genomes along with their metadata employed in this study.

**Supplementary Table 2**

MICs values for each drug tested on the 73 newly sequenced *A. baumannii* isolates of the HCG.

